# Increasing the Capping Efficiency of the Sindbis Virus nsP1 Protein Negatively Affects Viral Infection

**DOI:** 10.1101/453068

**Authors:** Autumn T. LaPointe, Joaquín Moreno-Contreras, Kevin J. Sokoloski

## Abstract

Alphaviruses are arthropod-borne RNA viruses that are capable of causing severe disease and are a significant burden to public health. Alphaviral replication results in the production of both capped and noncapped viral genomic RNAs, which are packaged into virions during the infections of vertebrate and invertebrate cells. However, the roles that the noncapped genomic RNAs (ncgRNAs) play during alphaviral infection have yet to be exhaustively characterized. Here, the importance of the ncgRNAs to alphaviral infection was assessed by using mutants of the nsP1 protein of Sindbis virus (SINV), which altered the synthesis of the ncgRNAs during infection by modulating the protein’s capping efficiency. Specifically, point mutants at residues Y286A and N376A decreased capping efficiency, while a point mutant at D355A increased the capping efficiency of the SINV genomic RNA during genuine viral infection. Viral growth kinetics were significantly reduced for the D355A mutant relative to wild type infection, whereas the Y286A and N376A mutants showed modest decreases in growth kinetics. Overall genomic translation and nonstructural protein accumulation was found to correlate with increases and decreases in capping efficiency. However, genomic, minus strand, and subgenomic viral RNA synthesis was largely unaffected by the modulation of alphaviral capping activity. In addition, translation of the subgenomic vRNA was found to be unimpacted by changes in capping efficiency. The mechanism by which decreased presence of ncgRNAs reduced viral growth kinetics was through the impaired production of viral particles. Collectively, these data illustrate the importance of ncgRNAs to viral infection and suggests that they play in integral role in the production of viral progeny.

**Importance:** Alphaviruses have been the cause of both localized outbreaks and large epidemics of severe disease. Currently, there are no strategies or vaccines which are either safe or effective for preventing alphaviral infection or treating alphaviral disease. This deficit of viable therapeutics highlights the need to better understand the mechanisms behind alphaviral infection in order to develop novel antiviral strategies for alphaviral disease. In particular, this study details a previously uncharacterized aspect of the alphaviral life cycle, the importance of noncapped genomic viral RNAs to alphaviral infection. This offers new insights into the mechanisms of alphaviral replication and the impact of the noncapped genomic RNAs on viral packaging.

## Introduction

Alphaviruses are positive sense, single-stranded RNA viruses which are capable of being transmitted between sylvatic vertebrate reservoir hosts and competent mosquito vectors in an enzootic cycle. During epizootic events, these viruses are capable of causing severe disease and are broadly categorized into two groups based on symptomology. Disease caused by the encephalitic alphaviruses, such as Venezuelan Equine Encephalitis virus, are characterized by neurological symptoms and are capable of causing severe encephalitis, which may result in the death of the vertebrate host (1, 2). In contrast, the arthritogenic alphaviruses, such as Sindbis virus (SINV), Ross River virus (RRV), and Chikungunya (CHIKV), cause disease ranging in severity from mild febrile illness to debilitating polyarthritis, which may persist for several months to years past the acute phase of infection (3−7). Despite the burden to public health posed by the alphaviruses, there are currently no effective and safe antiviral treatments for alphaviral disease.

During alphaviral infection, three viral RNA species are produced: the genomic strand, which encodes the nonstructural polyprotein, which is processed proteolytically to form the replication machinery; a minus strand RNA template; and a subgenomic RNA, which encodes the structural proteins. The positive sense alphaviral RNAs (vRNAs), namely the genomic and subgenomic vRNAs, have a type-0 cap structure co-transcriptionally added to their 5’ ends during RNA synthesis in order to protect the 5’ end of the transcript and to facilitate translation after entry to the host cell (8−10). The addition of the 5’ cap structure to the positive sense vRNAs is a multistep process involving at least two alphaviral nonstructural proteins. Briefly, the methyltransferase domain of the alphaviral nsP1 protein catalyzes the addition of a methyl group from S-andenosylmethionine to a GTP molecule, forming a covalent m^7^GMP-nsP1 intermediate (11). Following the removal of the 5’ γ-phosphate on the nascent vRNA molecule by nsP2, the m^7^GMP moiety is then transferred to the vRNA molecule by the guanylyltransferase activities of nsP1, resulting in the formation of the type-0 cap structure, ^7me^GpppA (12, 13). Importantly, biochemical studies of alphaviral nsP1 proteins have found that both the methyltransferase and guanylyltransferase activity of nsP1 can be altered either negatively or positively via point mutations *in vitro* (14−16).

The presence of the type-0 cap structure on the alphaviral genomic and subgenomic vRNAs were first observed by paper electrophoresis, or thin-layer chromatography, of metabolically labeled vRNAs that were chemically and enzymatically degraded (8, 9, 17). These studies further identified the sequences of the 5’ termini of the alphaviral genomic and subgenomic vRNAs, which provided the first evidence for independent promoter initiation for the two positive-sense vRNA species (9). While these studies were able to identify the presence of the 5’ type-0 cap structure, they were, by the nature of their design and technological limitations, unable to determine the relative frequency with which the positive sense vRNAs were capped.

Recently, we reported findings that indicated that the alphaviral genomic vRNAs are not ubiquitously capped, and that a significant proportion of the genomic vRNAs produced during SINV and RRV infection lack the 5’ type-0 cap structure (18). Furthermore, analyses of infectious and noninfectious viral particles demonstrated that both the capped and noncapped genomic vRNAs are packaged into viral particles throughout the course of infection. Through the use of tissue culture models of alphaviral infection, the presence of the noncapped vRNAs were found to correlate with the activation of a type-I IFN response. Moreover, an attenuated RRV mutant was found to produce fewer noncapped genomic RNAs relative to wild type virulent RRV (18, 19). Collectively, these data were highly suggestive of an important biological role for the noncapped genomic vRNAs during infection; however, the precise functions of the noncapped genomic vRNAs during infection remained unknown.

The goal of this study was to determine the importance of noncapped vRNAs to SINV infection of tissue culture cells. Here, we show that the capping activity of SINV nsP1 is capable of being modulated, both positively and negatively, via the mutation of specific residues within the Iceberg Domain, a structurally organized domain of cryptic function that is conserved amongst many viral capping enzymes (12). In addition, SINV infection was determined to be more detrimentally impacted by increasing the capping efficiency of SINV nsP1, rather than by decreasing efficiency. Specifically, increasing the rate of alphaviral capping negatively impacts viral growth kinetics by negatively affecting viral particle production. Collectively, our findings indicate that the noncapped vRNAs do play an important role during viral infection, and that decreasing the presence of noncapped vRNA impacts the viral life cycle at a fundamental level.

## Materials and Methods

### Tissue Culture Cells

BHK-21 cells (a gift from R.W. Hardy, Indiana University-Bloomington) were maintained in Minimal Essential Media (MEM, Cellgro) containing 10% fetal bovine serum (FBS, Atlanta Biologicals), 1% antibiotic-antimycotic solution (Cellgro), 1% nonessential amino acids (Cellgro), and 1% L-glutamine (Cellgro). Cells were cultured at 37^°^C and at 5% CO_2_ in a humidified incubator. Low passage stocks were maintained via regular passaging using standard subculturing techniques.

### Generation of SINV Capping Mutants

The SINV nsP1 mutants used in this study were generated either by site directed mutagenesis, or Gibson assembly reactions. Particularly, the SINV p389 Y286A and D355A mutants were generated via site-directed mutagenesis according to the instructions for Q5 site-directed mutagenesis kit (NEB). To this end, the parental wild type strain consisting of the p389 SINV nsP3-GFP reporter strain was PCR amplified with high fidelity Q5 DNA polymerase using primer sets that incorporated the indicated alanine substitutions, individually. Due to technical limitations that prevented the successful use of site-directed mutagenesis, the N376A mutant was generated by Gibson assembly via the Gibson Ultra kit (SGI) through the use of restriction digested p389 cDNA plasmid and a synthetic DNA fragment, according to the manufacturers’ instructions. SINV nanoluciferase reporters mutants, similar to those previously described, were generated by either site-directed mutagenesis, or by conventional restriction enzyme cloning schemes which swapped the GFP coding region of the existing p389 site mutants with a Nanoluciferase coding region in a modular fashion (18).

In all cases, individual mutants were verified by whole genome sequencing before proceeding to subsequent analyses. Full genome sequences are available upon request. Highly similar phenotypes were observed for any given mutant in all virus backgrounds evaluated.

### Production of SINV Wild Type and Mutant Virus Stocks

Wild type, Y286A, D355A, and N376A SINV p389 (a derivative of the Toto1101 strain containing GFP in frame with nsP3 (20)), as well as SINV pToto1101-nanoluc (a derivative of the Toto1101 strain containing nanoluciferase in frame with nsP3 (18)), were prepared by electroporation as previously described (21, 22). Briefly, 2.8×10^6^ BHK-21 cells were electroporated with 10μg of *in vitro* transcribed RNA using a single pulse at 1.5kV, 25mA, and 200Ohms from a Gene Pulser Xcell system (BioRad) as previously described (22). Cells were then incubated at normal incubation conditions until cytopathic effect became apparent, at which point the supernatant was collected and clarified by centrifugation at 8,000xg for 10min at 4°C. The clarified supernatant was then aliquoted into small-volume stocks, which were then stored at −80°C for later use.

### Analysis of Viral Growth Kinetics

To determine if mutation of the SINV nsp1 protein negatively affected infection, the viral growth kinetics of each of the individual strains described above were assayed in tissue culture models of infection. Essentially, BHK-21 cells were seeded in a 24-well plate and incubated under normal conditions. Once cell monolayers were 80-90% confluent, they were infected with either the wild-type parental virus, or the individual capping mutant viruses, at an MOI of 5 infectious units (IU) per cell. After a 1hr adsorption period, the inoculum was removed and the cells were washed twice with 1x PBS to remove unbound viral particles. Whole media was added, and the cells were incubated at 37^⁰^C in a humidified incubator in the presence of 5% CO_2_. At the indicated times post infection, tissue culture supernatants were harvested (and the media replaced), and viral titer was determined via plaque assay.

### Quantification of Infectious Virus by Plaque Assay

Standard virological plaque assays were used to determine the infectious titer of all viral samples produced during these studies. Briefly, BHK-21 cells were seeded in a 24-well plate and incubated under normal conditions. Once the cell monolayers were 80-90% confluent, they were inoculated with 10-fold serial dilutions of virus-containing samples. After a 1hr adsorption period, cells were overlaid with a solution of 0.5% Avicel (FMC Corporation) in 1x media for 28-30hrs (23). The monolayers were fixed with formaldehyde solution (3.8% formaldehyde in 1xPBS) for a period of no less than one hour. Plaques were visualized via staining with crystal violet after removal of the overlay and quantified by manual counting.

### XRN1 Protection Assay/ RppH Assay

Viral genomic RNAs were extracted from purified viral particles harvested at 24hpi. After extraction, the RNA samples were subjected to enzymatic degradation via the 5’→3’ exonuclease XRN-1, which is capable of degrading 5’ monophosphate and RNAs *in vitro*, but is unable to effectively degrade RNA substrates that are 5’ capped (24, 25). Briefly, equivalent amounts of viral genomic RNAs were incubated in the presence of 0.25 units of XRN-1 in a final volume of 20μl for a period of 5 minutes at 37° prior to being quenched via the addition of High Salt Column Buffer (25mM Tris-HCl, pH 7.6 / 400mM NaCl / 0.1% w/v Sodium Dodecyl Sulfate (SDS). After the termination of the reaction, the RNAs were purified via phenol:chloroform extraction and ethanol precipitation with linear acrylamide carrier. The resulting precipitate was resuspended and utilized as the substrate for the synthesis of cDNA via ProtoScript II reverse transcriptase (NEB). The resulting cDNAs were quantified via qPCR of an amplicon located in the nsP1 region, as described below, to determine the sensitivity of a sample relative to wild type viral genomic vRNAs (22, 26).

To confirm that the nature of the 5’ end, in particular the presence of a 5’ cap structure, was responsible for the differences in XRN-1 sensitivity the extracted RNAs were co-incubated with XRN-1 in the presence of the decapping enzyme RppH (18, 27, 28). To this end, the above reactions were supplemented with 1.25 units of RppH and processed as described above.

### Metabolic Labeling of Protein Synthesis

To determine the rates of host and viral during infection, BHK-21 cells were seeded in a 12-well tissue culture dish and grown to 80-90% confluence prior to being infected with either wild type parental virus, or one of the individual capping mutant viruses at an MOI of 10 IU/cell. After a 1hr adsorption period, fresh media was added to each well, and the cells were incubated under their normal incubation conditions. Thirty minutes before the indicated times post infection, the media was removed and replaced with methionine‐ and cysteine-free DMEM (Cellgro) to starve the cells of methionine. After a 30 minute incubation period, the starvation media was removed and replaced with methionine‐ and cysteine-free DMEM supplemented with 50μM L-azidohomoalanine (L-AHA), a methionine analogue (29, 30). After 2hrs, the labeling media was removed, the cells were washed with 1x PBS, and then harvested with RIPA buffer (50mM Tris-HCl, pH 7.5 / 150mM NaCl / 1% v/v Nonidet P-40 (NP-40) / 0.5% w/v SDS / 0.05% w/v Sodium Deoxycolate / 1mM EDTA). Cell lysates were labelled with DIBO-Alexa 648 (Invitrogen) at a final concentration of 5μM and incubated at room temperature in the dark for at least 1 hour. The labeled lysates were then loaded onto a 12% SDS-PAGE gel, and the proteins were separated by electrophoresis. Fluorescence was then imaged using a Pharos FX Plus Molecular Imager (BioRad), and densitometry of individual protein species was used to quantify viral and host protein expression.

### Western Blot Detection of SINV nsP2 Protein Expression

Whole cell lysates were generated from BHK-21 cells infected with either wild type, or individual SINV nsP1 capping mutant viruses by resuspension in RIPA buffer at 12 and 16hpi. Equivalent amounts of protein were loaded onto 10% acrylamide gels, and the individual protein species were resolved using standard SDS-PAGE practices. After sufficient resolution of the protein species, the proteins were transferred to PVDF membranes, which were rinsed in methanol and thoroughly dried after transfer. After drying the blots were probed with anti-SINV nsP2 polyclonal sera (a gift from R.W. Hardy at Indiana University-Bloomington), or anti-Actin (clone mAbGEa, ThermoFisher Scientific) antibodies diluted in 1xPBS supplemented with 1.0% v/v Tween (PBST) for a period of at least one hour at 25ºC under gentle rocking. The blots were washed three times with 1xPBST, and probed with fluorescent anti-rabbit (A32732, ThermoFisher Scientific) and anti-mouse secondary (A32723, ThermoFisher Scientific) antibodies. Protein detection was achieved using a Pharos FX Plus Molecular Imager (BioRad), and densitometry of individual protein species was used to quantify viral and protein expression.

### qRT-PCR Detection of SINV vRNAs

The quantitative detection of the SINV vRNAs was accomplished as previously described, with minor modifications (21). Briefly, to quantify the SINV genomic, subgenomic, and negative sense vRNAs, BHK-21 cells were infected at an MOI of 5 IU/cell and cells were harvested at 2, 4, and 8hpi via the addition of Ribozol (VWR). Total RNA was then isolated via extraction, and 1μg was reverse transcribed using Protoscript II Reverse Transcriptase (NEB) and a cocktail of specific RT primers based on the intended amplification targets. To detect the positive-sense RNA species the RT primer cocktail consisted of nsP1, E1, and 18S reverse primers; and to detect the minus strand the RT cocktail consisted of nsP1 forward and 18S reverse primers. The individual vRNA species were detected via TaqMan probes using the following primer sets: SINV nsP1 F 5’-AAGGATCTCCGGACCGTA-3’, SINV nsP1 R 5’‐ AACATGAACTGGGTGGTGTCGAAG-3’; SINV E1 F 5’‐ TCAGATGCACCACTGGTCTCAACA-3’, SINV E1 R 5’‐ ATTGACCTTCGCGGTCGGATACAT-3’; Mammalian 18S F 5’‐ CGCGGTTCTATTTTGTTGGT-3’, Mammalian 18S R 5’‐ AGTCGGCATCGTTTATGGTC-3’. The sequences of the TaqMan detection probes are as follows: SINV nsP1 Probe 5’-ACCATCGCTCTGCTTTCACAACGA-3’, SINV E1 Probe 5’‐ ACTTATTCAGCAGACTTCGGCGGG-3’, and Mammalian 18S Probe 5’‐ AAGACGGACCAGAGCGAAAGCAT-3’.

To quantify the total number of viral particles present in a sample, BHK-21 cells were infected at an MOI of 5 IU/cell and supernatant was collected at 24hpi. As previously described, 5uL of supernatant was reverse transcribed (21). Reverse transcription and qPCR reactions were performed identical to those above with the exception of the mammalian 18S primer and probes not being used. Absolute quantities of viral genomic RNAs were determined via the use of a standard curve of known concentration.

### Cell Viability Assay

To determine the effect of SINV infection on cell viability, BHK-21 cells were seeded in a 96-well plate and were infected with the wild type parental virus or the individual capping mutant viruses at an MOI of 10 IU/cell. After a 1hr adsorption period, whole medium was added and the cells were incubated under normal incubation conditions for the described times. Afterwards, the cells washed with 1x PBS and a solution of 1/6^th^ CellTiter 96 AQ_ueous_ One Solution Reagent (Promega) in 1x PBS was added to each well. The cells were then allowed to incubate at 37oC and 5% CO_2_ for 2hrs. Absorbance at 490nm was then recorded using a Synergy H1 microplate reader (BioTek).

### Quantification of Viral Gene Expression via Nanoluciferase Assays

To quantify genomic translation during infection, BHK-21 cells were infected at an MOI of 5 IU/cell with the wild type parental virus or the individual capping mutants containing the nanoluciferase gene within the nsP3 protein (18). After a 1hr adsorption period on ice, the inoculum was removed and the medium was replaced with pre-warmed whole medium. The infected cells were incubated under normal conditions, and at the indicated times post-infection, the medium was removed and the tissue culture cells were washed with 1x PBS. The cells were then harvested into a crude lysate by the addition of 1x PBS supplemented with 0.5% (v/v) Triton X-100. The lysate was transferred to a microfuge tube and frozen until the completion of the experimental time course. The samples were thawed and clarified by centrifugation at 16,000Xg for 5 minutes, and equal cell volumes of the nanoluciferase samples were processed using the Nano-Glo nanoluciferase assay system (Promega) according to the manufacturer’s instructions. Luminescence was then recorded using a Synergy H1 microplate reader (BioTek).

### Statistical Analysis

The quantitative data reported in this study are the means of a minimum of three independent biological replicates, unless specifically noted otherwise. The growth curve data presented in Fig.2 were statistically assessed using an area under the curve approach to determine differences in viral growth kinetics throughout the course of the assay. The statistical analysis of comparative samples was performed using variable bootstrapping where appropriate, as previously described (21). The error bars indicate the standard deviation of the mean. The *P* values associated with individual quantitative data sets are the result of Student’s *t* test for the corresponding quantitative data.

## Results

### SINV vRNA Capping Can be Modulated by Point Mutations in the nsP1 Protein

As described above, we recently reported that a significant number of SINV genomic vRNAs produced during infection lack the type-0 5’ cap structure. Despite being noninfectious, the noncapped viral genomic RNAs (ncgRNAs) are produced in a temporally dependent manner and are packaged into viral particles throughout the course of infection (18). Given the evolutionary conservation of the production of the ncgRNAs during alphaviral infection, we hypothesized that they must be biologically important to viral infection. This led us to question how modulating capping activity, which would alter the production of capped viral genomic RNAs and ncgRNAs, impacts viral infection. In order to determine the biological importance of the ncgRNAs during viral infection, we modulated the methyltransferase and guanylyltransferase activities of the alphaviral nsP1 protein to generate mutant SINV strains with either increased or decreased rates of viral capping. These mutations were based on previous work done in VEEV by Li et al. (2015), where alanine substitutions at specific residues affected the nsP1 protein’s methyltransferase and guanylyltransferase activities, as well as cap formation as a whole. Specifically, the previous study found that alanine substitutions at Y286 and N376 decreased RNA capping efficiency, although to different extents, whereas an alanine substitution at D355 increased RNA cap formation. All three residues are highly conserved across multiple alphaviruses, including the model alphavirus SINV (Fig.1A). Modeling of the SINV nsP1 protein using ITASSER indicates that residues D355 and N376 are likely close to one another, in parallel alpha helices proximal to the catalytic site within the Iceberg Domain (Fig. 1B)(31). Nonetheless, Y286 is located at a distal site far from the putative active sites, as identified by prior mutational analyses (11, 12, 16). Admittedly, the ITASSER predicted structure is unlikely to be a wholly accurate representation of the true structure of the SINV nsP1 protein. Despite the inherent inaccuracy of protein folding prediction algorithms in the absence of a closely related crystal structure, secondary and tertiary structural elements reminiscent of methyltransferase and guanylyltransferase enzymes may be detected, including a Rossmann fold-like structure surrounding the key residues involved in methyltransferase activity (16, 32). In addition, the overall arrangement of the Core Region and Iceberg Domains are largely intact, and the known membrane anchoring domains face outward from the globular body of the protein (32). The differential ability of the Y286, D355, and N376 to affect VEEV nsP1 capping activity as well as their locations in highly conserved domains, which play integral roles in cap formation, made them strong candidates for testing how modulating the capping activity of nsP1 affects SINV infection.

**Figure 1.**
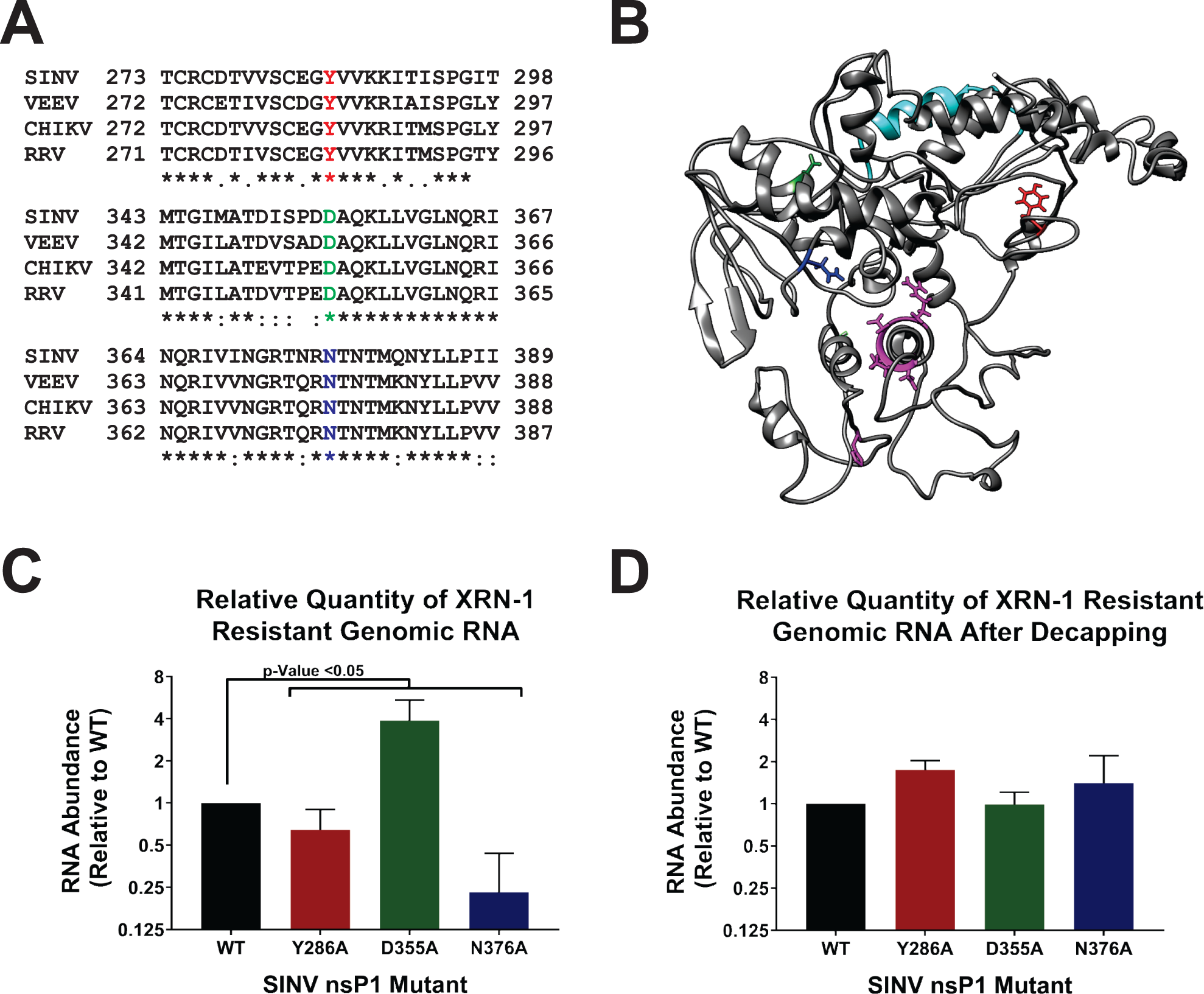
Point Mutations In The Nsp1 Protein Of SINV Alter 5’ vRNA Capping Efficiency. (A) Amino acid sequence alignment of selected alphavirus nsP1 proteins. The individual nsP1 protein sequences of Sindbis (ViPR-U38305), Venezuelan Equine Encephalitis (VEEV, ViPR-L01443), Chikungunya virus (CHIKV, ViPR-DQ443544), and Ross River virus (RRV, ViPR-GQ433359) were aligned by Clustal Omega. (B) An ITASSER predicted structure of the SINV nsP1 protein. Amino acid residues of importance are highlighted as follows-Red=Y286, Green=D355, Blue=N376, Cyan=amphipathic helix, and the residues involved in the methyltransferase activities, including H39 which binds to the ^m7^GMP residue are pink. (C) A graph depicting the relative quantity of XRN-1 resistant vRNA isolated from viral particles produced by BKH-21 cells infected with wild-type SINV or the individual capping mutants 24hpi. (D) Identical to panel C, with the exception that the viral genomic RNAs were enzymatically decapped concurrent with XRN-1 treatment. All quantitative data shown represents the means of a minimum of three independent biological replicates utilizing 3 independent particle preparations, with the error bars indicating the standard deviation of the means. P-Values, as indicated on the figure, were determined by Student’s T-test.

In order to verify that the SINV nsP1 mutations altered the rate of vRNA capping, we assessed what proportion of viral particles contained capped RNA genomes. We have shown previously that both capped genomic RNAs and ncgRNAs are packaged and released during viral infection (18). Thus, knowing what proportion of capped viral RNA is being packaged into viral particles gives insight into the general capping efficiency of nsP1 during viral infection. To this end, the comparative assessment of viral capping was accomplished by comparing the relative sensitivity of the mutants to RNAse degradation using XRN-1, a 5’-3’ exoribonuclease which preferentially degrades RNAs lacking a protective 5’ cap structure, in particular 5’ monophosphate RNAs (24, 25). By measuring the amount of RNA resistant to XRN-1 degradation, we were able to determine what relative proportion of the total input RNA was capped relative to wild type SINV, as the predominant noncapped 5’ end was previously determined to be a 5’ monophosphate (18). As shown in Figure 1C, both SINV N376A and SINV Y286A had significantly lower proportions of XRN-1 resistant genomic RNAs compared to WT, with SINV Y286A having approximately 25% less XRN-1 resistant RNA compared to WT, and SINV N376A having approximately 75% less XRN-1 resistant RNA. In contrast, and as expected from the previous study by Li et al. (2015), SINV D355A was found to have a significantly greater proportion of XRN1-resistant RNA, with a mean approximately four-fold greater than what was observed for WT SINV. To confirm that the observed resistance of the SINV D355A genomic RNAs was due to a cap structure, the genomic RNAs were subjected to XRN-1 nuclease treatment in the presence of an established decapping enzyme (18, 27, 28). As shown in Figure 1D, enzymatic removal of the 5’ cap structure eliminated the resistance of the D355A derived genomic RNAs to XRN-1 treatment, resulting in XRN-1 resistances comparable to WT SINV.

From these data, we were able to conclude that the point mutations engineered into the SINV nsP1 protein can indeed alter capping activity, both negatively, in the case of Y286A and N376A; and positively, as with D355A. Therefore, the individual nsP1 mutants can be used to assess the impact of the ncgRNAs by modulating the capping activity of nsP1 in tissue culture models of infection in a controlled manner.

### Increased Capping Decreases SINV Growth Kinetics in Mammalian Cells

While the effects of synonymous point mutations on nsP1 capping efficiency have been previously characterized for VEEV at an enzymatic level (14), the effects of modulating capping activity on viral infection as a whole has not yet been studied in detail. Given the molecular function of the 5’ cap structure, one could expect that the SINV nsP1 mutants that decreased vRNA capping would show impaired viral growth kinetics relative to wild type virus. Likewise, if the ncgRNAs were truly nonfunctional, a mutant that increased capping would show enhanced viral growth kinetics in regards to wild type infection. As demonstrated by the data presented in Fig. 2A, decreasing SINV capping modestly decreased viral growth kinetics, as observed for the SINV Y286A and SINV N376A mutants. However, the SINV D355A mutant, which increased vRNA capping relative to WT, exhibited significantly decreased titers over the course of infection (Fig.2A). In addition to a 2-log decrease in viral titer, the SINV D355A mutant produced plaques approximately half the size of those produced by wild type SINV (Fig.2B and 2C). Despite not showing significantly decreased viral growth kinetics, both the SINV Y286A and SINV N376A mutants also exhibited a small plaque phenotype, albeit not to the same extent as the SINV D355A mutant. In addition, all three capping mutants were found to have decreased induction of cell death compared to WT SINV (Fig.2D).

**Figure 2.**
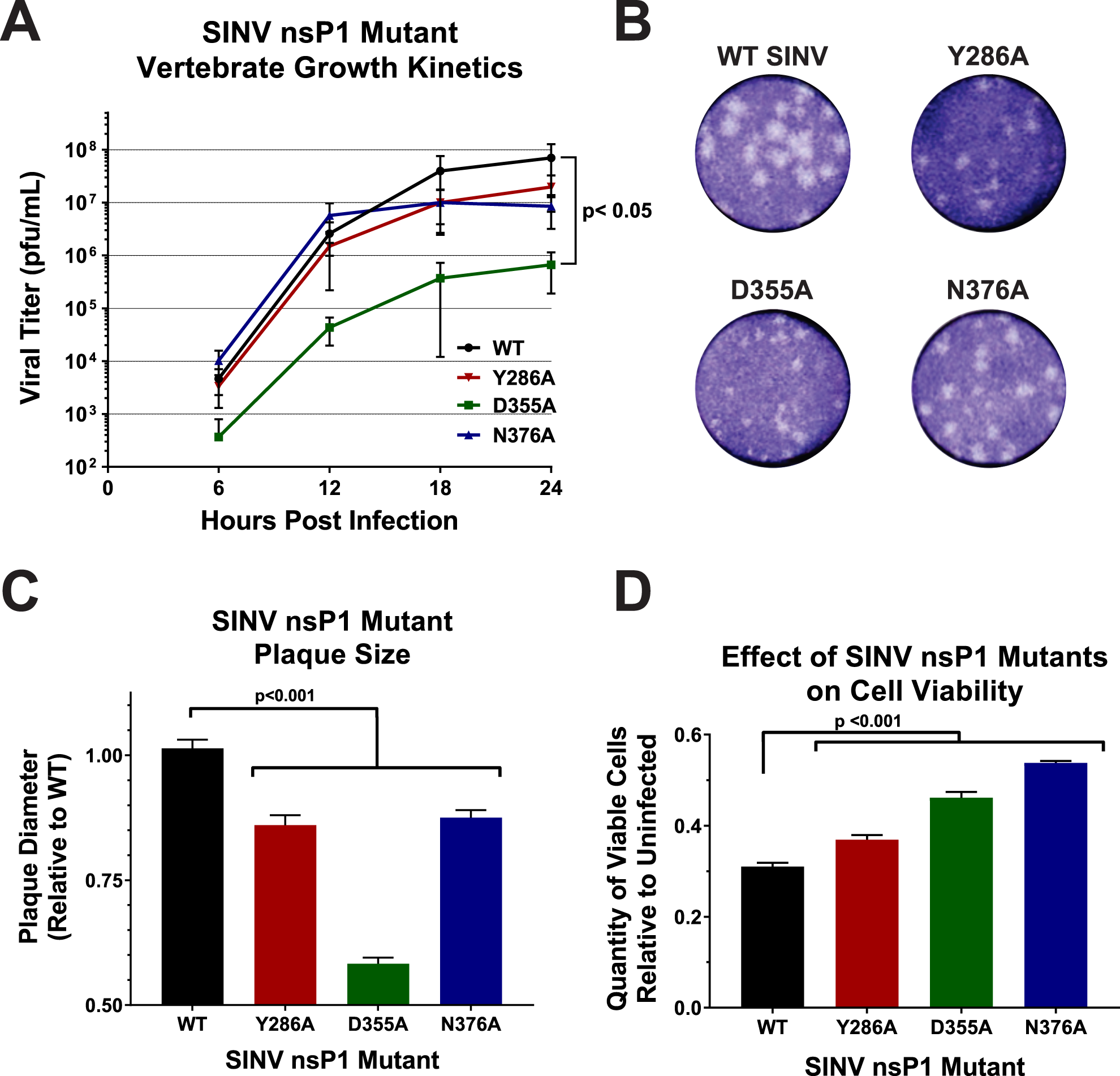
Altering Viral Capping Efficiency Negatively Impacts Viral Infection. (A) One-step growth kinetics of the individual capping mutants and parental wild type SINV as observed in BHK-21 cells infected at an MOI of 0.5 PFU/cell. Statistical significance was determined by area under the curve analysis. (B) Plaque morphology of wild type SINV and capping mutant viruses in BHK-21 cells overlaid with 1 0.5% solution of Avicel at 24h post-infection. (C) A graph indicating the average plaque diameter of the mutant viruses relative to wild type SINV. Size was determined by ImageJ software (NIH). (D) Cell viability of BHK-21 cells infected with the individual capping mutants at 24hpi relative to mock infected BHK-21 cells. All quantitative data shown represent the means of at least three independent biological replicates, with the error bars representing standard deviations of the mean. Statistical significance, as indicated within each panel, was determined by Student’s *t* test.

Collectively, these data indicate that changing the efficiency of vRNA capping negatively impacted viral growth kinetics, illustrating that a step in the viral life cycle has been detrimentally affected. Furthermore, the viral growth kinetics data suggests that SINV is more sensitive to increased vRNA capping efficiency than it is to decreased capping efficiency. Thus, the ncgRNAs produced during infection are indeed biologically important. Nevertheless, despite the clear negative impact to viral infection, the precise nature of the molecular consequences of increased vRNA capping and decreased presence of ncgRNAs cannot be determined from these data alone. As such, the viral gene expression and vRNA synthesis profiles of each of the SINV nsP1 mutants were next assessed to determine if they were negatively affected by modulation of vRNA capping.

### Translation of the SINV Genomic RNA Correlates with Capping Efficiency

For the majority of mRNAs, a key factor for determining whether or not an mRNA is translated is the presence of a functional 5’ cap structure (33−35). Therefore, changing the ncgRNAs to capped genomic RNAs or vice versa by altering viral capping efficiency should impact the amount of protein being produced by viral RNA during infection. To investigate the effect(s) that modulating nsP1 capping efficiency has on viral gene expression, a SINV reporter, which contained the open reading frame of nanoluciferase in frame with the nsP3 nonstructural protein (18), was used to measure viral genomic RNA translation throughout infection (Fig.3A). Nanoluciferase expression was measured at regular intervals over the course of viral infection (Fig. 3B). As expected, the SINV D355A mutant, which increased capping relative to wild type, exhibited increased genomic vRNA translation compared to WT SINV for every time point measured (Fig. 3C). Similarly, SINV N376A, which decreased capping relative to wild type, showed decreased translation (Fig. 3C). Curiously, the SINV Y286A mutant showed differential nanoluciferase expression during infection, with translation being slightly increased very early during infection, decreased at 8hpi, and similar to wild type at later times post infection.

**Figure 3.**
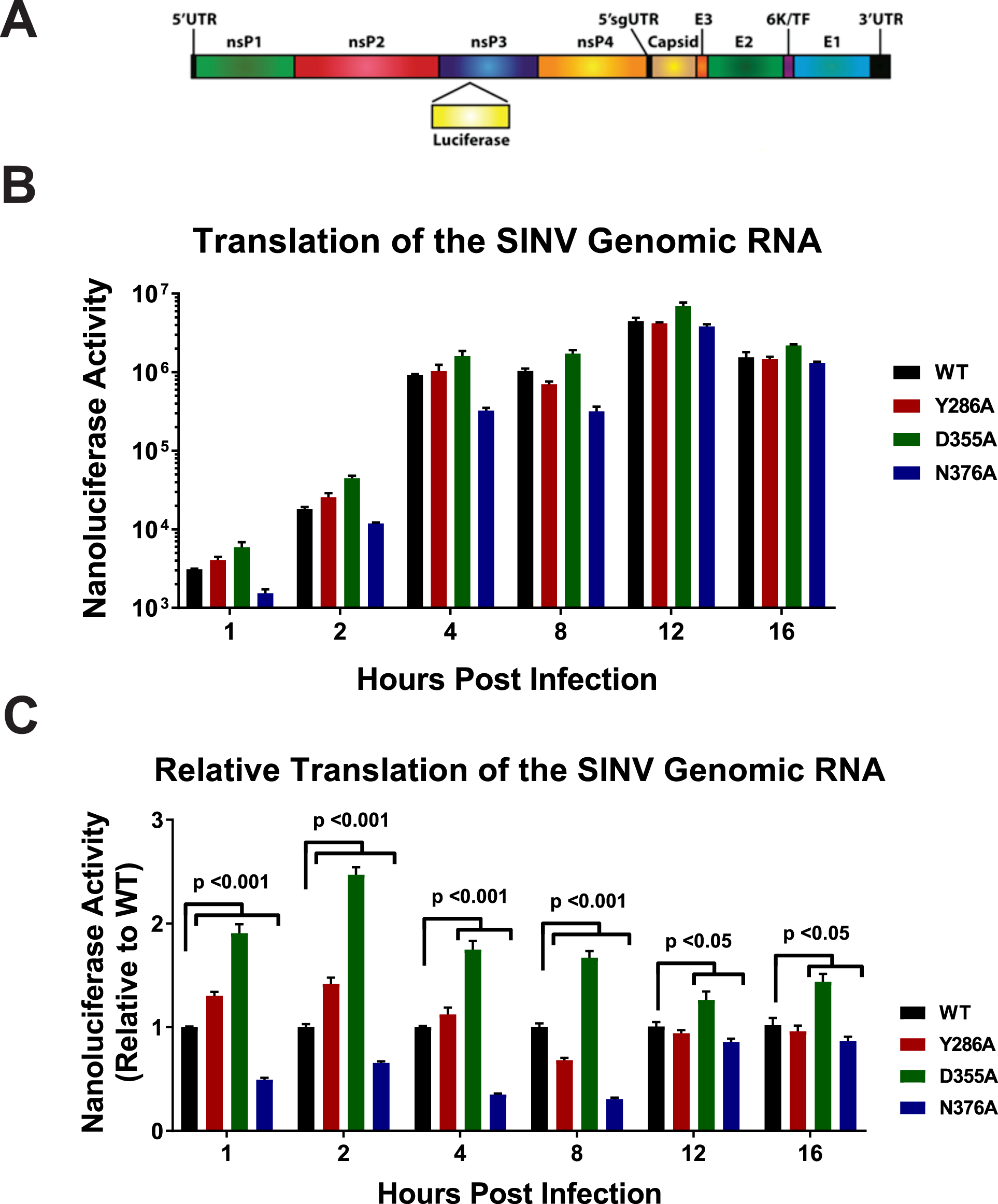
Translation of the Genomic vRNA Correlates with Viral Capping Efficiency. (A) Schematic diagram of the SINV nanoluciferase reporter used in this study. (B) BHK-21 cells were infected with either parental wild type, or an individual SINV capping mutant nanoluciferase reporter strain. The level of nanoluciferase activity was quantified at the indicated times post infection. C) The nanoluciferase activity, as reported in panel B, normalized to wild type expression at each individual time point to enable readers to identify differences in translation. All the quantitative data shown represent the means of three independent biological replicates, with the error bars representing standard deviations of the mean. Statistical significances, as indicated in the figure, were first determined using ANOVA analyses followed by post-hoc statistical analyses by Student’s *t* test.

While the SINV nanoluciferase reporter virus allows for the accurate quantification of viral gene expression during infection, it does not measure the sum accumulation of the nonstructural proteins. To measure the relative accumulations of the viral nonstructural proteins, the levels of SINV nsP2 protein were assessed via Western blotting at 8, 12 and 16hpi. As shown in Fig. 4A and quantified in. Fig 4B, at 8hpi the accumulation of the SINV nsP2 proteins between wild type SINV, SINV D355A, and SINV N376A differed to a statistically significant extent. The abundance of nsP2 was increased ~2.5-fold during SINV D355A infection, relative to wild type SINV. In contrast, nsP2 levels during SINV N376A infections were decreased ~2-fold relative to wild type SINV infections. The SINV Y86A mutant was more or less equivalent to wild type nsP2 levels. Similar trends were observed at later times during infection. As depicted in Figs. 4C and 4D, at 12hpi the SINV nsP2 levels were statistically increased relative to wild type infection; however, given the low magnitude of effect, these differences are unlikely to be biologically significant. In contrast, at 16hpi SINV nsP2 levels were increased to an extent that is likely biologically meaningful (Figs 4E and 4F).

**Figure 4.**
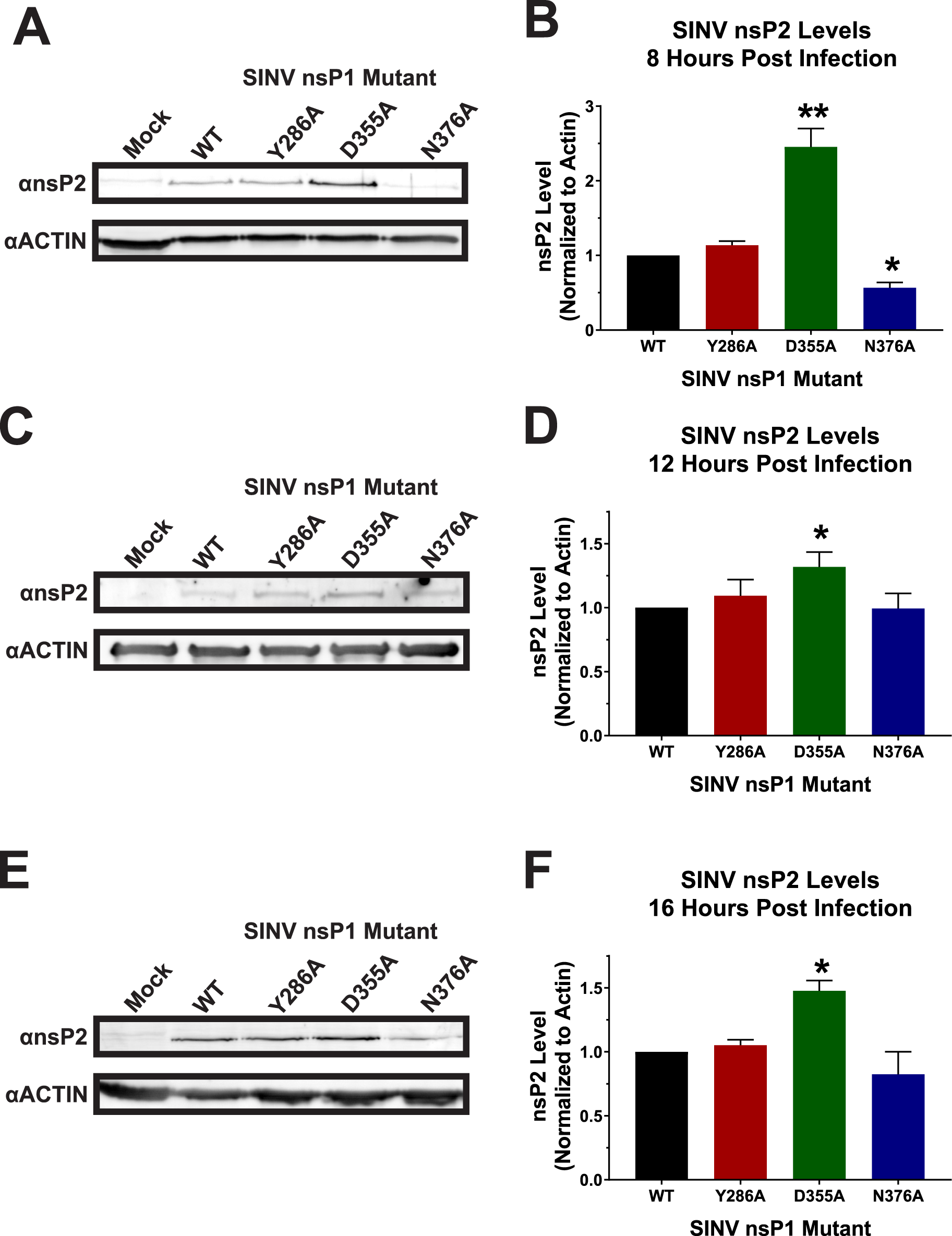
nsP2 Protein Levels are Impacted by Mutation of the nsP1 Protein. (A) BKH-21 cells were infected with wild type SINV or an individual capping mutant at an MOI of 5 PFU/cell and assessed by western blotting to determine the abundance of nsP2 at 8hpi. Actin is shown as a loading control. B) Densitometric quantification of the nsP2 protein normalized to Actin levels at 8hpi. Panels C and D) Western blots and densitometry analyses identical to those described for panels A and B, with the exception that the timing of the assay coincided to 12hpi. Panels E and F) Western blots and densitometry analyses identical to those described for panels A and B, with the exception that the timing of the assay coincided to 16hpi. The Western blot images shown are representative of at least three independent biological replicates. All the quantitative data shown represent the means of three independent biological replicates, with the error bars representing standard deviations of the mean. Statistical significances, as indicated in the figure, were first determined using ANOVA analyses followed by post-hoc statistical analyses by Student’s *t* test.

Overall, the levels of genomic translation for the SINV nsP1 mutants nicely reflect their relative differences in capping efficiency, with increased vRNA capping resulting in increased translation of the nonstructural proteins, and decreased vRNA capping resulting in decreased translation at 8hpi. These data are consistent with the conversion of the ncgRNAs to translationally competent capped genomic vRNAs. Furthermore, western blotting confirmed the differences in translational activity detected during SINV nanoluciferase reporter infections. Taken together, the nanoluciferase and western blot data indicate that the translation of the genomic vRNAs continues during infection with SINV D355A beyond what is observed for wild type SINV. Nonetheless, the biological differences in translational activity observed between the SINV D355A mutant and its respective parental wild type strain fail to explain the approximately 200-fold reduction in viral titer.

### Modulating SINV Capping Does Not Alter Overall RNA Synthesis or Accumulation

Since differences in viral gene expression failed to outright explain the observed decreased viral titers associated with the SINV D355A mutant, we next sought to identify if vRNA synthesis was impaired as a result of nsP1 mutation. To determine what impact altering viral capping efficiency has on vRNA synthesis, the genomic, subgenomic, and minus strand vRNAs were quantitatively assessed at 2, 4, and 8 hours post infection via qRT-PCR. At 2hpi, the SINV D355A mutant, which has increased vRNA capping relative to wild type SINV, produced slightly more SINV genomic and subgenomic RNAs; whereas the SINV N376A mutant, which exhibits decreased vRNA capping, produced slightly fewer genomic and subgenomic RNAs (Fig. 5A). In general, the differences in the amount of genomic and subgenomic RNAs being produced by these two capping mutants is reflective of the differences observed in the synthesis of their replication machinery (Figs. 3 and 4). At 4hpi, all three SINV nsP1 mutants show a statistically significant deficit in the amount of subgenomic RNA present (Fig. 5B). Nonetheless, these deficits are unlikely to be biologically meaningful due to their overall magnitude of effect relative to wild type SINV. By 8hpi however, all three capping mutants show similar levels of all three RNA species compared to WT, with the exception of N376A, which had less minus strand template compared to WT (Fig. 5C).

**Figure 5.**
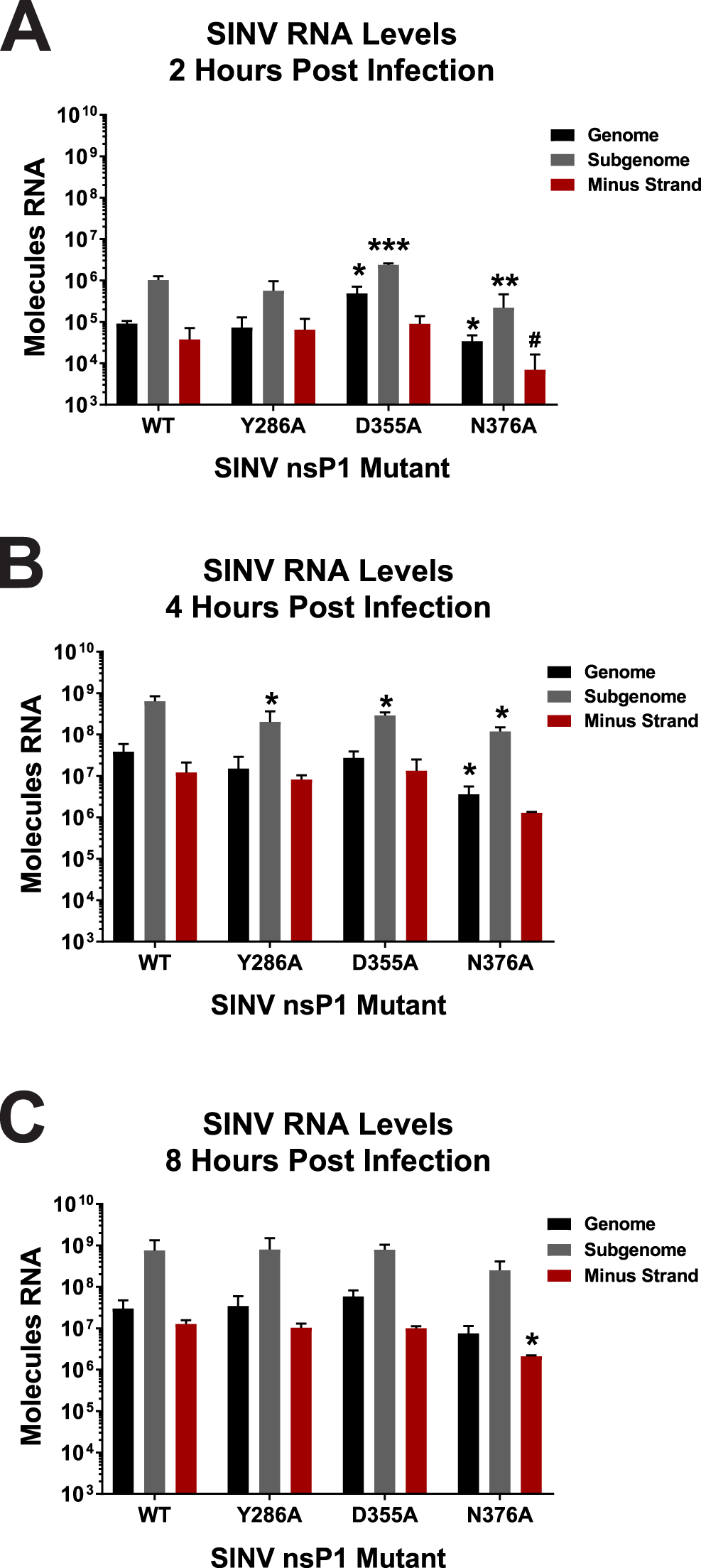
Altering vRNA Capping Efficiency Impacts Early RNA Synthesis. (A) BHK-21 cells were infected with either wild type parental SINV or an individual capping mutant virus at an MOI of 5 PFU/cell. At 2 hours post infection, the total cellular RNA was extracted and assessed for the absolute quantities of the genomic, subgenomic, and minus-strand vRNAs by qRT-PCR. (B and C) Identical to panel A with the exceptions that the time points are 4 and 8 hours post infection, respectively. All the quantitative data shown represent the means of three independent biological replicates with the error bars representing standard deviations of the mean. Statistical significances, as indicated in the figure, were first determined using ANOVA analyses followed by post-hoc statistical analyses by Student’s *t* test. P-Values as determined by Student’s *t* test are represented by *, p<0.05. **, p<0.01. ***, p<0.001. #, one of the biological replicates was below limit of detection precluding meaningful statistical analysis.

Altogether, these data demonstrate that altering the capping activity of nsP1 may have impacts on RNA synthesis very early during viral infection, but that these differences become muted as infection progresses. Importantly, these data indicate that vRNA synthesis on the whole has not been disrupted, precluding the possibility that the point mutations made in nsP1 have disrupted the function of the other nonstructural proteins. Moreover, similar to that described above regarding viral gene expression in the previous section, the ~2-log reduction in viral growth kinetics observed for the SINV D355A mutant is not due to decreased vRNA accumulation or defective vRNA synthesis.

### Translation of the SINV Subgenomic RNA is Largely Unaffected by Viral Capping

Given that neither the differences in nonstructural gene expression or vRNA synthesis were capable of explaining the negative impact of the SINV nsP1 D355A mutation, we next sought to determine if the function of the subgenomic vRNAs were impacted. As both the genomic and subgenomic vRNAs are capped by nsP1, it was hypothesized that mutations which increased capping of the genomic vRNAs might impact subgenomic vRNA function. As such, it could be expected that these vRNAs would exhibit similar responses to altered capping in terms of translation. In order to determine whether translation of the subgenomic vRNAs were impacted similarly to the genomic vRNAs, the amount of protein produced late during infection was measured using metabolic labeling (Fig. 6A). Curiously, both the SINV D355A and SINV N376A mutants exhibited a modest decrease in viral capsid production (Fig. 6B). In addition, none of the SINV nsP1 mutants exhibited differences in regards to the shutoff of host translation, a hallmark of alphaviral infection as reviewed in (36), as evidenced by the labeling of the cellular Actin protein, relative to WT SINV (Fig. 6C). Interestingly, however, the ongoing synthesis of a high molecular weight protein consistent with the nonstructural polyprotein was reproducibly detected during the metabolic labeling of SINV D355A mutant infections.

**Figure 6.**
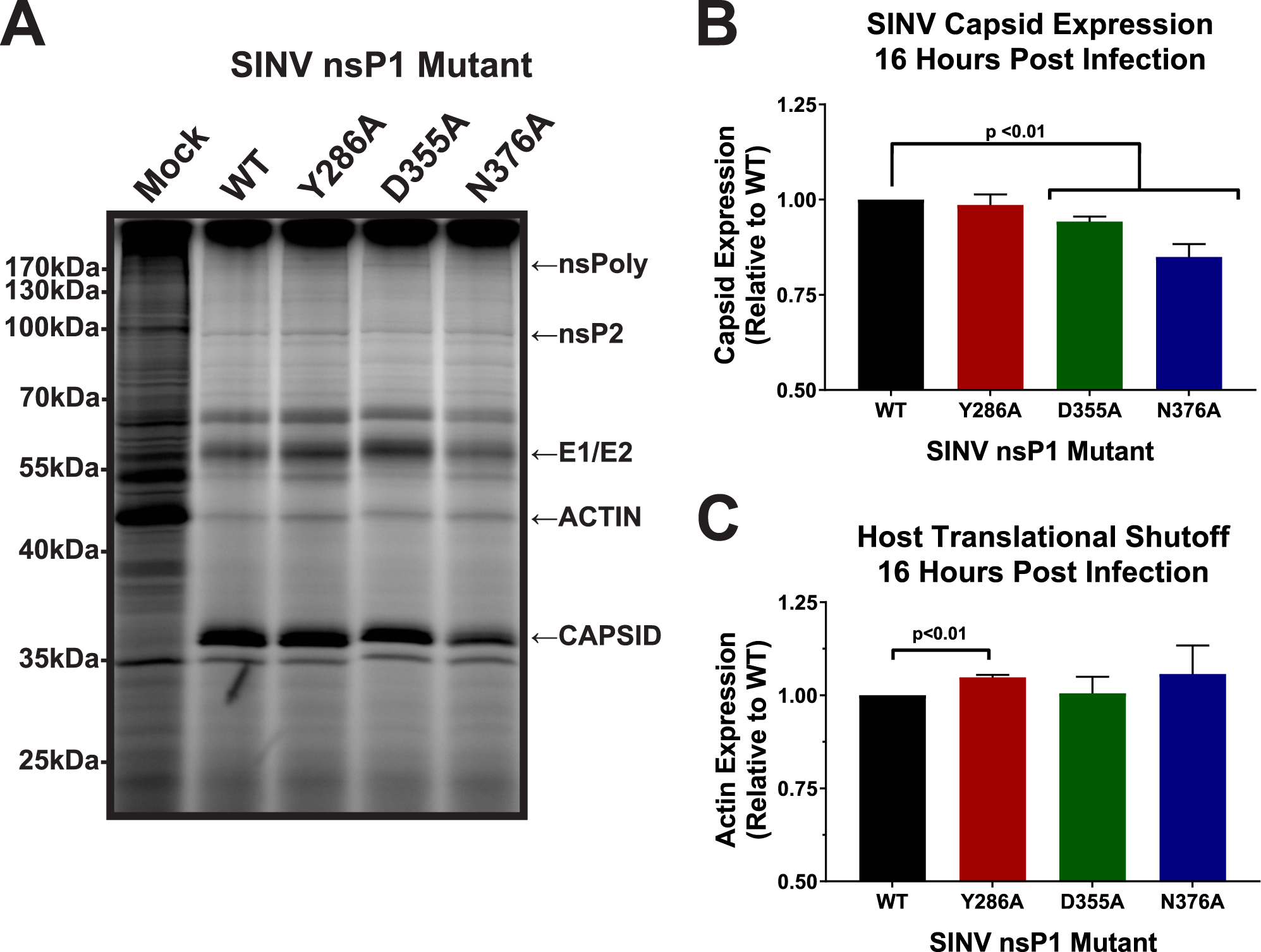
Subgenomic Gene Expression Is Unaffected by Altering SINV vRNA Capping. (A) BHK-21 cells were either mock treated or infected with wild type SINV or an individual capping mutant at an MOI of 10 PFU/cell. At 14hpi, the cells were pulsed with L-AHA for a period of 2 hours. Afterwards, the cells were harvested and equal cell volumes of cell lysate were analyzed by SDS-PAGE and fluorescent imaging. The data shown are representative of three independent biological replicates. (B) Densitometric quantification of the SINV capsid protein, with intensity relative to wild-type SINV shown. (C) Densitometric quantification of the host actin protein with intensity relative to wild type SINV. All the quantitative data shown represents the means of three independent biological replicates with the error bars representing standard deviations of the mean. Statistical significance was determined by Student’s *t* test.

Collectively, these data indicate that changes in capping efficiency of the genome may either not lead to equivalent changes in capping of the subgenomic vRNA; or alternatively, that SINV, and likely other alphaviruses, regulate the translation of the structural proteins differently than the nonstructural proteins (37). Regardless, these data suggest that subgenomic translation and host translational shutoff are not impacted to a biologically significant degree by modulating the efficiency of vRNA capping, and thus are not responsible for the decreased viral titer associated with increased capping efficiency.

### Increasing SINV Capping Decreases Viral Particle Production

Given that the molecular characterizations of the SINV nsP1 mutants had so far failed to identify the molecular defect leading to decreased viral growth kinetics, we expanded our analyses beyond the lifecycle events most obviously affected by the 5’ cap structure. Since vRNA synthesis and viral gene expression were unaffected, we next sought to determine whether viral particle assembly was negatively impacted by increasing the capping efficiency of the SINV nsP1 protein.

To determine whether increasing capping efficiency affected viral particle production, we quantified the total number of particles produced by each mutant as well as wild type SINV after 24hrs of infection. Similar to what was observed during the kinetic analyses of viral infection, increased vRNA capping was associated with the production of significantly fewer particles, with the SINV D355A mutant producing ~25-fold fewer particles (as measured by genome equivalents per ml) than wild type SINV (Fig. 7A). In contrast, particle production was largely unaffected during infections of the SINV Y286A and SINV N376A mutants, which decreased vRNA capping.

**Figure 7.**
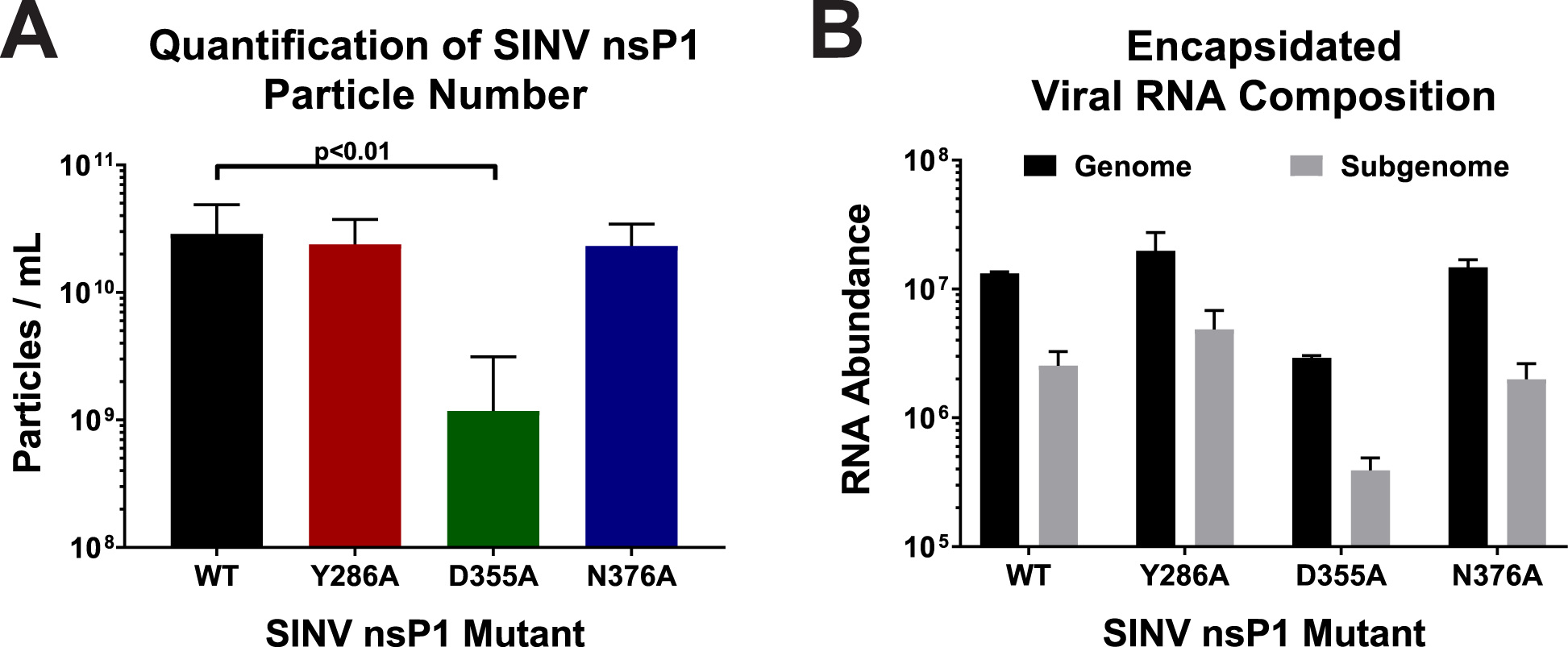
Analysis of SINV Particle Production. A) BHK-21 cells were infected with either wild-type SINV or an individual capping mutant at an MOI of 5 PFU/cell. At 24hpi the total number of viral particles produced was measured using qRT-PCR. Data shown represents the means of at least 6 independent biological samples. B) Quantitative determination of the composition of the encapsidated viral RNAs in mature extracellular viral particles. Samples of virus containing supernatants were assessed to determine the absolute quantities of the genomic and subgenomic RNAs via standard curve qRT-PCR. All the quantitative data shown represent the means of three independent biological replicates, with the error bars representing standard deviations of the mean. Statistical significances, as indicated in the figure, were first determined using ANOVA analyses followed by post-hoc statistical analyses by Student’s *t* test.

The point mutants utilized in this study are located within, or closely adjacent to, the SINV packaging signal (38). Previous characterizations of the alphaviral packaging signals have indicated the importance of stem-loop structures which contained a guanosine triplet in the loop region. Mutational analyses of the alphaviral packaging elements determined that mutation of the guanosine triplets, or deletion of the packaging element altogether, significantly reduced viral titer due to nonselective particle assembly leading to the production of alphaviral particles containing the subgenomic vRNAs (38). Further experiments defined that a minimum of two guanosine triplet stem loops was sufficient to impart wild type particle production with selectivity for the genomic RNA. Even though none of the point mutants used in this study impact either the guanosine triplets, or the general secondary structures (as predicted by *in silico* analysis), the potential for the inhibition of assembly by a non-capping mechanism existed, warranting further assessment of particle production during SINV nsP1 mutant infections. As mentioned earlier, disruption of the alphaviral packaging signal leads to the production of viral particles containing predominantly subgenomic RNAs (38). As shown in Fig. 7B, quantitative determinations to identify which specific vRNAs were packaged into wild type and SINV nsP1 mutant viral particles indicated no aberrant packaging of the SINV subgenomic RNA, consistent with an intact functional packaging signal.

The high degree of similarity between the relative magnitude of effect regarding decreased viral titer and viral particle production during SINV D355A infection indicates that particle assembly, and not the slight perturbations in vRNA synthesis or structural gene expression, is primarily responsible for decreased viral growth kinetics. Importantly, these data suggest that SINV, on the whole, is much more tolerant of mutations which decrease capping efficiency than those that increase capping efficiency, and that by increasing the capped to noncapped viral RNA ratio, viral particle production has been detrimentally affected. Moreover, these data validate that the packaging signal remains functional in the presence of the nsP1 point mutations described in this study.

## Discussion / Conclusion

### The capping of SINV genomic vRNAs can be modulated by point mutations in nsP1

The data shown in Fig. 1 indicates that, in SINV, 5’ capping of the genomic vRNA can be modulated via single point mutations in the nsP1 protein. This is true for both increasing vRNA capping, as seen with SINV D355A, and decreasing capping, as seen with SINV Y286A and SINV N376A. Moreover, vRNA capping can also be modulated to different extents, as seen with SINV Y286A and SINV N376A, which decreased vRNA capping by 25% and 75%, respectively. The ability to change nsP1 capping efficiency in a controllable manner opened up new avenues to explore the molecular and biological importance of both the capped genomic vRNAs and the ncgRNAs during infection in tissue culture models of infection and *in vivo*.

From the data above, we may conclude that a primary consequence of mutating the SINV nsP1 protein is the alteration of vRNA capping. Nonetheless, the alphaviral nonstructural proteins interact with one another during infection (39−42). Previous studies have shown that disrupting these interactions by mutation results in poorer viral infection and, more specifically, leads to severe defects in vRNA synthesis (39−41). However, the disruption of nonstructural protein interactions is unlikely with the nsP1 mutants reported in this study, as the phenotypes described for situations where the interactions between the nonstructural proteins have been disrupted are inconsistent with what is reported here. For example, several residues in nsP4 have been reported as important for interactions with nsP1, such as G38L in nsP4 (39). When this residue in nsP4 was mutated, viral infection exhibited decreased growth kinetics and a small plaque phenotype. However, a hallmark of disrupting the nonstructural protein interactions was the severely decreased synthesis of minus strand vRNA throughout the course of infection (39, 41). As seen in Fig. 5, none of the nsP1 mutants examined in this study exhibited a significant deficit in production of any viral RNA species, with the exception of a minor decrease in minus strand vRNA synthesis by N376A. It is of note that the nsP4 G38L mutant was able to be rescued by an additional mutation in nsP1 at N374 (39). The nsP1 N374 mutation resulted in complete restoration of viral titer and partial restoration of minus strand vRNA synthesis in the nsP4 G83L background compared to wild type. While the effect of mutating nsP1 N374 on capping efficiency is not known, one could speculate that, given its proximity to other residues which we have shown alter capping activity, that modulating the capping efficiency of nsP1 could be a way of coping with the detrimental nsP4 G38L mutation, which creates a severe defect in RNA synthesis. In addition to this, a previous study has reported that the region in nsP1 encompassing the SINV point mutations utilized here possess little to no interaction with the nsP2 protein (42). Thus, for the reasons described above, the nsP1 residues mutated during this study are likely not involved in mediating nonstructural protein interactions, and the resulting deficits in the viral lifecycle are not due to disrupted nonstructural protein interactions.

### SINV infection is more sensitive to increased capping than decreased capping

Multiple studies have shown that polymorphisms in nsP1 have profound effects on virulence. Mutations in regions of nsP1 have been shown to alter vRNA synthesis, viral titer, viral sensitivity to IFN, and disease severity *in* vivo (19, 39, 43). Certain residues, such as nsP1 H39 in SINV, completely abrogate viral infection by eliminating the methyltransferase activity of the nsP1 protein (15, 16, 44). Therefore, we expected that incorporating the point mutations which alter viral capping would have impacts on viral infection. However, we were surprised to find that while the increasing capping mutant was found to have decreased viral growth kinetics, the decreased capping mutants were not significantly different from wild type SINV (Fig. 2A). This result was especially surprising when it was found that all three capping mutants had small plaque phenotypes and decreased cell death, yet D355A was the only mutant to show altered growth kinetics (Fig. 2B, C, and D). Serial passaging of the mutants used in this study have indicated that they are stable for at least 4 sub-passages, as no reversion events (based on plaque phenotype) were observed for any of the SINV nsP1 mutants. This suggests that SINV is more detrimentally impacted by changes which increase the amount of capped vRNA present than those which decrease capping efficiency. Thus, increasing the capping efficiency of the SINV nsP1 protein, which effectively reduced the production of the ncgRNAs, indicates that the ncgRNAs are biologically important to viral infection.

### Altering capping leads to changes in genomic translation but not RNA synthesis or subgenomic translation

As would be expected, changes in nsP1 capping efficiency correlated with changes in genomic vRNA translation. In addition, the reduced presence of the ncgRNAs correlated with increased translation of the viral genomic RNA throughout infection, as can be seen with the increased capping mutant SINV D355A. Furthermore, decreasing the capping efficiency of the nsP1 protein, as evidenced by the SINV N376A mutant, modestly decreased translation. Admittedly, the second decreased capping mutant, SINV Y286A, did not follow the same pattern as the SINV N376A mutant. This may be due to the comparatively minor decrease in capping efficiency caused by SINV Y286A not being significant enough to consistently alter the vRNA population leading to dysregulated genomic translation.

However, the increased translation exhibited by the SINV D355A mutant did not lead to lasting compounding biological effects, at least in tissue culture models of infection. For instance, despite there being more replication machinery being produced early during SINV D355A infection, there were no overt differences in the synthesis of any of the viral RNA species during the time points tested (Fig. 5). This suggests that increasing or decreasing the production of the nonstructural proteins alone is not enough to alter RNA synthesis over the long term in highly permissive tissue culture models of infection. Nonetheless, due to technical limitations we were unable to accurately assess vRNA synthesis earlier than two hours post infection. Hence, the possibility that vRNA synthesis is enhanced very early during infection remains unaddressed.

Another unexpected result was that changes in capping efficiency seems to affect subgenomic translation differently than genomic translation. The data presented in Fig. 6 show that both the increased and decreased capping mutants SINV D355A and SINV N376A demonstrate slight, biologically unmeaningful, decreases in capsid production at 16hpi, despite having notable differences in nonstructural gene expression (Figs. 3 and 4). This suggests that either capping efficiency is regulated differently for the subgenomic RNA than for the genomic RNA, resulting in no differences in capping efficiency for the subgenomic RNA, or that translation of the subgenomic RNA is less dependent on the presence of a 5’ cap than the genomic RNA during infection. The latter is supported by previous studies which demonstrate that the eIF4F complex, which includes the cap-binding eIF4E, is not needed in order to initiate translation of the alphaviral subgenomic RNA (37, 45). These previous studies, along with the data presented here, suggests that translation of the subgenomic RNA is unaffected, at least in part, by modulation of vRNA capping.

In addition to there being little effect on subgenomic translation, there was also shown to be no differences in terms of host translational shutoff between any of the capping mutants and WT SINV (Fig. 6C). This further supports that the point mutations made in nsP1 are not negatively affecting interactions between nonstructural proteins, because disruptions between nsP1 and nsP4 have been previously found to negatively affect host translational shutoff (39). Host translational shutoff, for SINV at least, has been largely attributed to the translation of the structural proteins (46). Therefore, it is unsurprising that there is no change in host shutoff between the capping mutants and the WT SINV given that there is little difference in subgenomic expression.

### Decreased titer due to changes in capping caused by interference with particle production

The decreased viral growth kinetics observed during SINV D355A infection correlated remarkably with decreased particle production. However, the congruence of the decrease in infectious units and viral particles infers that viral infectivity is, more or less, identical for all of the viral strains. The discrepancy between the two magnitudes of effect (an approximate 5-fold difference) are likely due to confounding variations in the accuracy and precision of the two measurements. Nonetheless, it remains possible that the viral particles have differences in their basal infectivity. Studies examining earlier effects of the ncgRNAs are ongoing and will be presented in a follow up study.

The observation that decreased particle production is the primary molecular defect during SINV D355A infections suggests that increasing the amount of capped genomic RNAs, thereby decreasing the number of ncgRNAs, negatively impacts the assembly of nascent viral particles. Whether or not the increased capping activity is directly, or indirectly, responsible for the packaging phenotype isn’t definitively known. Characterizations of SINV packaging indicates selectivity remains intact during the SINV D355A assembly process despite decreased particle production overall, as the packaging of subgenomic vRNAs was not observed. This is indicative of a functional alphaviral packaging signal despite the incorporation of minor point mutants into the region defined as the packaging signal for SINV. Moreover, the SINV Y286A mutant which also resides within the SINV packaging element lacks an appreciable packaging phenotype. Thus, the assembly phenomena associated with the SINV D355A mutant cannot be simply explained by disruption of the alphaviral packaging signal.

While the underlying mechanism is unclear, the data presented above indicates that nonstructural protein expression is increased relative to wild type parental virus, and remains increased well into the late stages of infection during SINV D355A infections. Collectively, these data indicate that decreasing the production of the ncgRNAs perturbs viral genomic RNA function beyond the individual RNA level, as apparent compounding effects on the genomic RNA population are observed. Precisely how the translationally inactive ncgRNAs serve to modulate genomic vRNA function as a whole, leading to efficient particle assembly, is unknown.

We propose that, as diagrammed in Fig. 8, during wild type infections, the translationally inactive, ncgRNAs temper the molecular activities of the translationally active, capped genomic vRNA population allowing for the temporal progression of infection to lead to the assembly and release of viral particles. We postulate that the ncgRNAs, due to their lack of translational capacity, interact with a unique set of host factors relative to the capped translationally competent genomic RNAs. Collectively, these interactions lead to the development of a pro-assembly microenvironment by excluding host factors that either inhibit the assembly process, or promote nucleocapsid disassembly. For instance, if the ncgRNAs foster an non-translational environment through the interaction of host factors, such as those found within stress granules, the 60S ribosomal subunit, which is implicated in nucleocapsid disassembly would be excluded from the local microenvironment, allowing assembly to occur unimpeded (47, 48). However, when the genomic RNA population is altered by increasing the efficiency of genomic RNA capping, such as observed with the SINV D355A mutant, the increased and continuous translational activity of the genomic vRNA culminates in the formation of a pro-translational vRNA “pool” that is refractory to encapsidation and particle assembly. Work examining such possibilities are ongoing in the Sokoloski lab and will be reported in the future.

**Figure 8.**
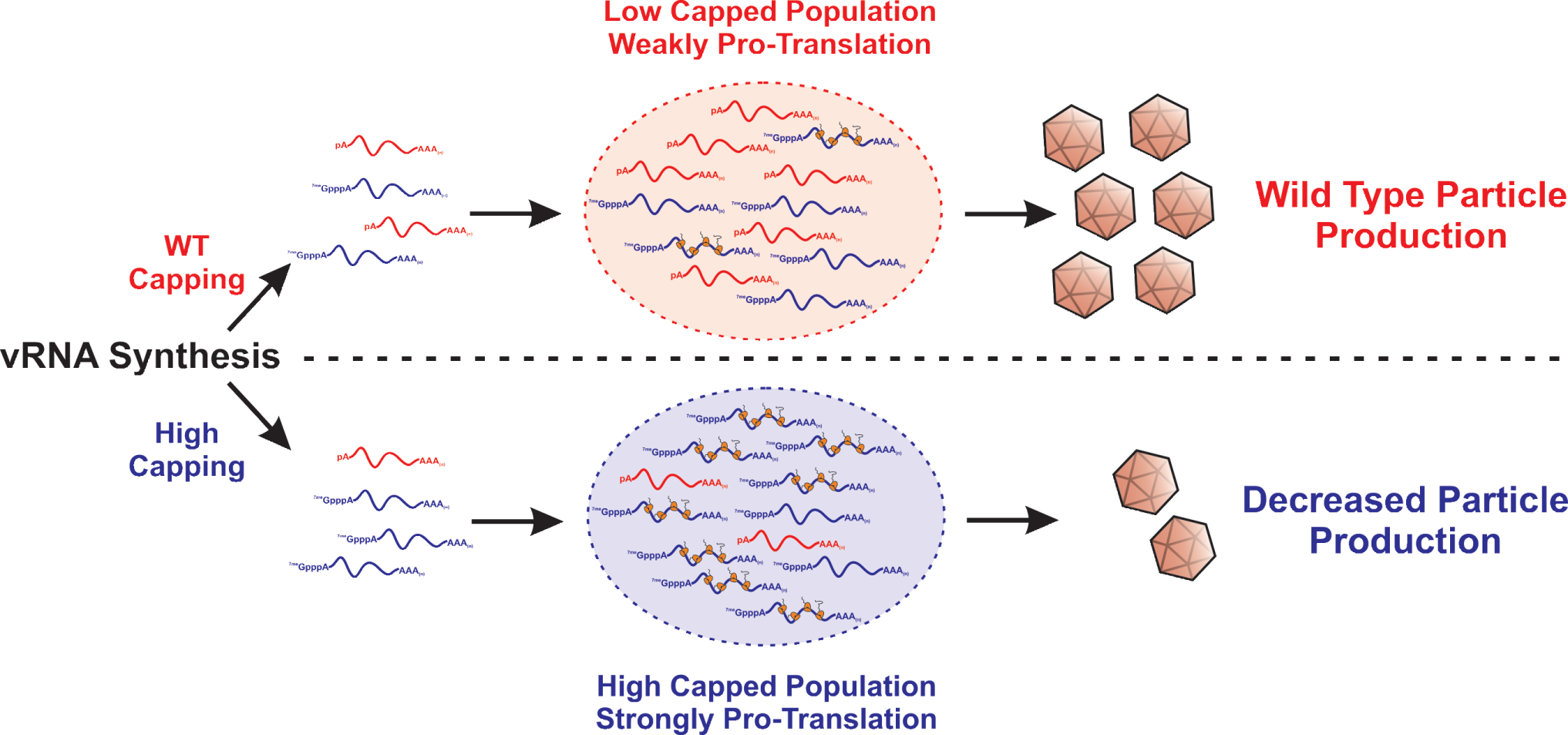
Proposed Model of How Increasing Genomic vRNA Capping Negatively Impacts Viral Infection.

## Conclusions

Collectively these data affirm the existence and biological importance of the ncgRNAs during SINV infection. This assertion is directly supported by the capacity to modulate the capping activities of nsP1 protein via site directed mutagenesis resulting in the increased production of XRN-1 resistant genomic vRNAs without increased overall genomic RNA numbers. Moreover, the preponderance of gene expression data indicating increased translation brought about by increasing the capping of the genomic vRNA supports the existence of the alphaviral ncgRNAs at a functional level. Finally, the molecular characterizations of SINV nsP1 mutant infections provides insight into the biological importance of the ncgRNAs in regards to the regulation of alphaviral infection at the molecular level.

## Acknowledgments

We would like to thank the members of the Sokoloski, Chung, and Lukashevich labs for their input during the development of the project and the preparation / editing of the manuscript. In addition, we would like to thank Dr. Susana Lopez for the opportunity to host Joaquín Moreno-Contreras during the course of this work.

